# An *Nlrp5*-null mutation leads to attenuated *de novo* methylation in oocytes, accompanied by a significant reduction in DNMT3L

**DOI:** 10.1101/2025.02.20.639225

**Authors:** Leah Nic Aodha, Alexandra Pokhilko, Leah U Rosen, Styliani Galatidou, Edyta Walewska, Christian Belton, Antonio Galvao, Hanneke Okkenhaug, Lu Yu, Asif Nakhuda, Bill Mansfield, Soumen Khan, David Oxley, Montserrat Barragán, Gavin Kelsey

## Abstract

*Nlrp5* encodes a core component of the subcortical maternal complex (SCMC) a cytoplasmic protein structure unique to the mammalian oocyte and cleavage-stage embryo. *NLRP5* mutations have been identified in patients presenting with early embryo arrest, recurrent molar pregnancies and imprinting disorders. Correct patterning of DNA methylation over imprinted domains during oogenesis is necessary for faithful imprinting of genes. It was previously shown that oocytes with mutation in the human SCMC gene *KHDC3L* had globally impaired methylation, indicating that integrity of the SCMC is essential for correct establishment of DNA methylation at imprinted regions. Here, we present a multi-omic analysis of an *Nlrp5*- null mouse model, which in GV oocytes displays a misregulation of a broad range of maternal proteins, including proteins involved in several key developmental processes. This misregulation likely underlies impaired oocyte developmental competence. Amongst impacted proteins are several epigenetic modifiers, including a substantial reduction in DNMT3L; we show that *de novo* DNA methylation is attenuated in *Nlrp5*-null oocytes. This provides evidence for mechanisms leading to downstream misregulation of imprinted genes, which in turn, may result in imprinting syndromes, multi-locus imprinting disturbances (MLID) and hydatidiform moles.

## Introduction

The mammalian oocyte provides not only the maternal genetic contribution to the embryonic genome, but also the proteins and mRNAs necessary to sustain the newly fertilised zygote through the first stages of embryogenesis, until zygotic genome activation and beyond^1–4^. During primordial germ cell specification, the epigenetic landscape is erased and re-established after sex specification, in order to restore gametic pluripotency. As the growing oocyte develops, DNA methylation patterning is re-established in a transcription-dependent manner, and is completed upon reaching the surrounded nucleolus (SN) stage of nuclear maturation^5^, at which point the oocyte becomes largely transcriptionally silent^6–9^. The DNA methylation landscape in the oocyte differs significantly from that of somatic cells. While considered a repressive modification in somatic cells, DNA methylation in oocytes does not coincide with silenced gene regulatory elements. Instead, DNA methylation is present in a bimodal pattern of hypermethylated but actively transcribed regions, and hypomethylated transcriptionally inactive regions^8^. Later stages of oocyte maturation are orchestrated by its pre-existing pool of mRNAs and proteins^10^. Upon fertilisation, the newly created zygote undergoes another wave of global DNA methylation erasure, with the exception of methylation at imprinted regions, which is retained to ensure parent-of-origin gene expression^7,11^.

While the establishment of DNA methylation in the mouse oocyte is catalysed by DNA methyltransferase DNMT3A and requires its cofactor DNMT3L^12,13^, the maintenance of methylation through cell divisions post-fertilisation is dependent on DNMT1, regulated by accessory proteins UHRF1 and DPPA3^14–17^. DNMT1 also maintains parent-of-origin specific methylation at imprinted regions during global methylation erasure events^18^. Maintenance post-fertilisation of methylation specifically at imprinted regions also requires the KRAB-zinc finger proteins ZFP57 and ZNF445 together with TRIM28 (KAP1, TIF1B) ^19,20^. Both UHRF1 and DNMT1 can also contribute to *de novo* methylation in oocytes^14^ ^21^ ^22^, and UHRF1 has been shown to play other important roles in the oocyte, such as protecting against DNA damage^22^, and regulating the cytoplasmic architecture and function in the oocyte via tubulins and other microtubule-related proteins^23^.

However, UHRF1 and DNMT1 are considered particularly crucial for maintenance of DNA methylation. DNMT3A and DNMT3L are predominantly localised to the nucleus in oocytes and cleavage-stage embryos, whereas DNMT1 and its accessory protein UHRF1 are predominantly localised to the cytoplasm during this developmental window^14^, although some DNMT1/UHRF1 is retained in the nucleus^18,24^, indicating an important role of regulated sub-cellular localisation in controlling DNA methylation dynamics. Given that later stages of oogenesis and early stages of embryogenesis are directed by the oocyte pool of proteins and mRNAs, investigating the dynamics of the oocyte proteome may hold the clues to understanding the mechanisms by which DNA methylation establishment and maintenance are regulated, with important implications for the understanding of DNA methylation-related disorders, such as multi-locus imprinting disturbances (MLIDs) and imprinting syndromes in humans^25^.

The subcortical maternal complex (SCMC) is an enigmatic protein complex unique to the mammalian oocyte and cleavage-stage embryo. This complex is composed of several of the most highly abundant oocyte proteins, encoded by maternal-effect genes, although its function remains to be fully characterised^26–28^. Structures of the core complex, comprising NALP5 (or NLRP5, MATER), OOEP (FLOPED), TLE6, and KHDC3 (FILIA), in both human and mouse, have recently been solved^29,30^.

Recent studies on SCMC proteins have focused on their relationship with cytoplasmic lattices (CPLs), another previously described oocyte structure with incomplete characterisation of function. CPLs are found throughout the cytoplasm, and are likely composed of the core SCMC and associated proteins including PADI6, suggesting that SCMC proteins are not in fact subcortical in nature^3,31–33^. In light of this new framing, SCMC-CPL structures are thought to be involved in sequestration and protection of important developmental proteins from premature degradation^3,31,34^. Interestingly, mutations in SCMC genes have been associated with aberrantly high DNA methylation in murine pre-implantation embryos, in some cases coupled with altered localisation of DNMT1 and UHRF1^35,36^. Mutant variants of SCMC genes such as *KHDC3L*, *PADI6* and *NLRP5* have been identified in patients presenting with recurrent biparental complete hydatidiform moles^37^, and offspring with MLID and imprinting disorders^38,39^, suggesting a link between the SCMC and either establishment or retention of DNA methylation patterning at imprinted genes.

Furthermore, human oocytes with inactivating mutations in *KHDC3L* are globally hypomethylated^37^. The difficulty in determining a causal link between SCMC variants and downstream disordered DNA methylation lies in the diversity of clinical outcomes of SCMC mutations in humans, coupled with limited human material for analysis, and the fact that SCMC-null mutations in mice usually lead to embryonic arrest prior to zygotic genome activation^34,40–43^. Even when human SCMC-related imprinting disorder variants are mimicked in the murine system they have been shown to lead to 2-cell arrest^35^. Nonetheless, the tractability of mouse knockouts enables them to be a valuable tool in analysing the consequences of inactivating SCMC mutations, despite potential imperfections.

We focus on understanding the potential causes of developmental incompetence in oocytes using a novel mutation in the core SCMC gene *Nlrp5.* To understand how disruption of the SCMC in oocytes might lead to disordered DNA methylation and adverse developmental outcomes, we employ a multi-omic approach, profiling the oocyte proteome, transcriptome, and DNA methylation landscape, in addition to investigating the localisation of key epigenetic modifier proteins in *Nlrp5 -/-* oocytes.

## Results

### NALP5 staining in *Nlrp5* mutant oocytes shows whole cytoplasm localisation

*Nlrp5* encodes the NALP5 protein, which when absent or severely depleted results in the SCMC losing its structural integrity^26^. Severe depletion of CPLs^32^, an increase in disordered mitochondrial localisation and activity^40^, and altered endoplasmic reticulum distribution and calcium homeostasis^44^ in murine oocytes have also been reported. *Nlrp5-*deficient oocytes from previous models lead to embryonic arrest at the early cleavage stages when fertilised by wild-type sperm^34,45^. To interrogate the consequences of SCMC disruption on oocyte and pre-implantation embryo development, we generated a new knockout mouse model with an inactivating deletion in the core SCMC gene *Nlrp5.* We employed CRISPR/Cas9 technology to create an *Nlrp5* knockout by inducing a double-stranded break (DSB) within the *Nlrp5* gene, which led to a 5-base pair (bp) deletion, on a C57BL/6Babr genetic background. Our deletion, hereby referred to as *Nlrp5 -*, produced a premature stop codon, causing the mRNA to be targeted for nonsense-mediated decay (Figure 1A). Ablation of the NALP5 protein was tested in *Nlrp5 +/-* and *Nlrp5 -/-* germinal vesical (GV) stage oocytes via confocal immunofluorescence imaging (Figure 1B). Immunofluorescence imaging demonstrated that not only is the NALP5 protein ablated in *Nlrp5 -/-* oocytes, but it is also reduced in *Nlrp5 +/-* oocytes, where the prominent subcortical signal for NALP5 observed in *Nlrp5 +/+* oocytes is lost, and becomes more evenly distributed throughout the cytoplasm. Although the term SCMC was coined because of the apparent immunofluorescence localisation of its constituent proteins to the oocyte subcortex, this observation, coupled with recent colocalization studies^3,32^, suggests that the observed subcortical localisation of these proteins is likely an artefact of oocyte permeabilization and staining protocols leading to oversaturation with secondary antibodies at the subcortex^33^. The fact that NALP5 is one of the most abundant proteins in the oocyte and a significant reduction in NALP5 in *Nlrp5 +/-* oocytes reduces this subcortical saturation and leads to a more evenly dispersed cytoplasmic staining supports recent studies that indicate presence of the SCMC throughout the oocyte cytoplasm^3^.

**Figure 1:**
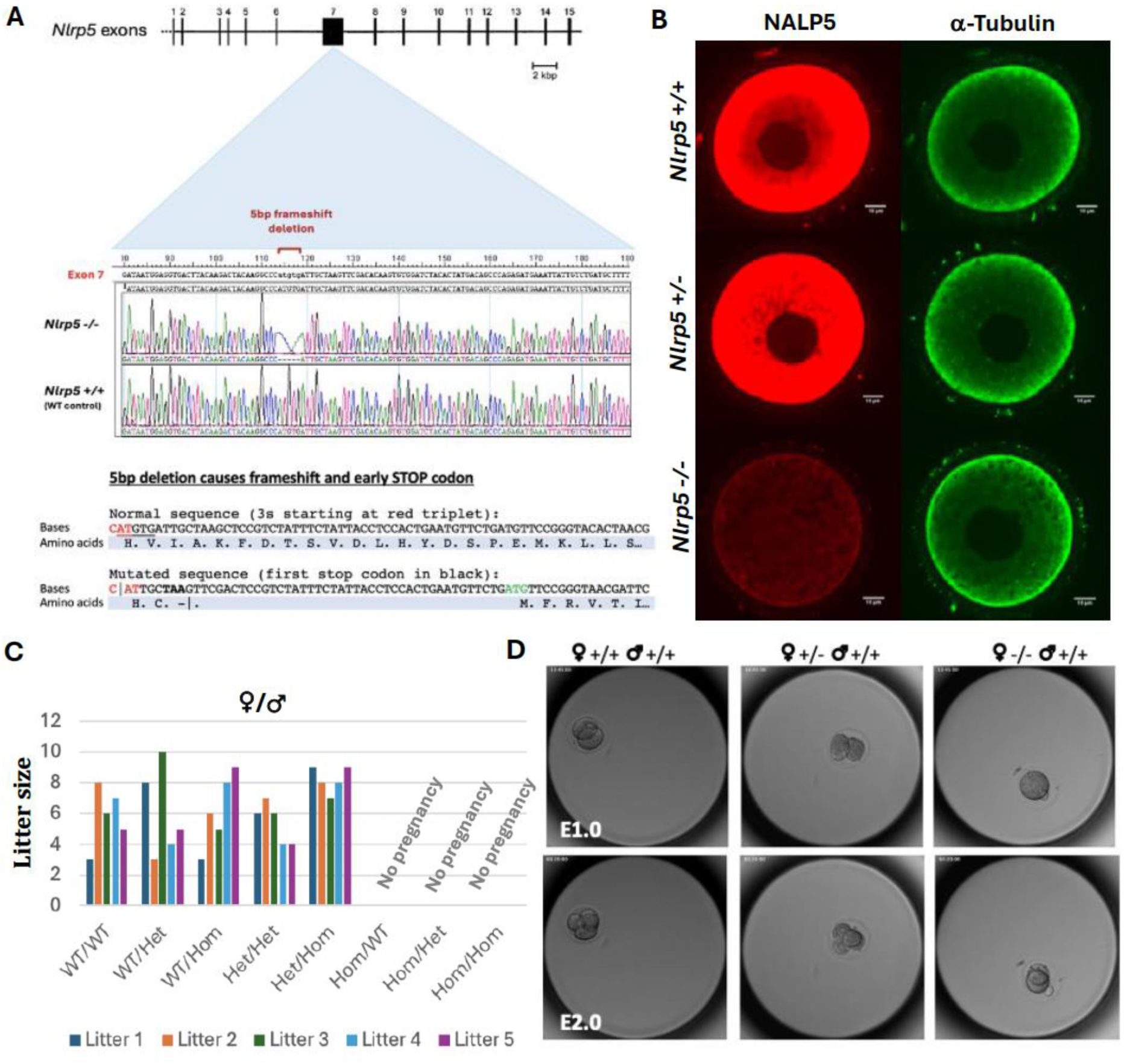
***Nlrp5* knockout mouse design and characterisation. (A)** Schematic of target sequence (base and amino acid mutation) for CRISPR/Cas9-mediated knockout generation, **(B)** Representative images showing immunofluorescence verification of NALP5 protein ablation (red) in germinal vesicle (GV) *Nlrp5 +/+, +/-* and *-/-* oocytes. ⍺-tubulin staining control (green). 10µm scalebar. n = 6 *Nlrp5 -/-*, 6 *Nlrp5 +/-,* 4 *Nlrp5 +/+*. **(C)** Clustered column chart showing number of pups per litter for mating pairs of each genotype combination. No pregnancies in matings with *Nlrp5 -/-* females. WT = *Nlrp5 +/+,* Het = *Nlrp5 +/-,* Hom = *Nlrp5 -/-.* **(D)** Representative time-lapse images of wild-type and *Nlrp5* mutant embryos at E1.0 and E2.0, demonstrating 2-cell arrest of maternal *Nlrp5* knockout-derived embryo (n = 16 ♀*Nlrp5 +/+*, 11 ♀*Nlrp5 +/-*, 16 ♀*Nlrp5 -/-)*.

### Impaired nuclear maturation and developmental competence of *Nlrp5 -/-* oocytes is accompanied by differences between *Nlrp5* -/- and *Nlrp5 +/+* oocyte transcriptomes

*Nlrp5 -/-* and *Nlrp5+/-* mice were phenotypically indistinguishable from *Nlrp5 +/+* mice in soma, though *Nlrp5 -/-* females were infertile, as previously reported^45^. Inter- genotype crosses set up to determine if mutants had differences in fertility or litter sizes showed that *Nlrp5* -/- males and *Nlrp5 +/-* mice of either sex had normal fertility, and litter sizes were not significantly different from wild-type pairings (Figure 1C). *In vitro* fertilisation (IVF) was performed on oocytes collected from 4-10-week-old *Nlrp5 -/-, Nlrp5 +/-,* and *Nlrp5 +/+* mice, fertilised with wild-type C57BL/6Babr sperm. Fertilised zygotes were cultured in a time-lapse imaging incubator for 72 hours, showing that our *Nlrp5* knockout recapitulated the 2-cell embryonic arrest phenotype reported in previous *Nlrp5* mutants^40,45^. Fertilised *Nlrp5 -/-* oocytes did not progress past the 2-cell stage of embryogenesis (Figure 1D), with most arresting or becoming visibly degenerated prior to the first cell division. In several cases, embryos that did achieve the 2-cell stage underwent reverse cleavage, before degenerating. *Nlrp5 +/-* oocyte-derived embryos developed as normal.

During the GV stage of oocyte maturation, chromatin undergoes a conformational change, moving from the less mature ‘non-surrounded nucleolus’ (NSN) stage, to the more mature surrounded nucleolus (SN) stage, at which point the oocyte becomes largely transcriptionally silent^46,47^. Although oocytes may bypass the NSN-SN transition during maturation, oocytes that have not successfully achieved the SN checkpoint typically lead to cleavage-stage embryonic arrest when fertilised^48^. Upon fluorescence imaging of Hoechst-stained GV oocytes, we observed that very few *Nlrp5 -/-* oocytes achieved the SN stage of oocyte nuclear maturation, and those that did had a qualitatively different SN conformation than *Nlrp5 +/-* and *Nlrp5 +/+* oocytes (Figure 2A, B, C). This nuclear maturation impairment was detectable in oocytes from *Nlrp5 -/-* females as young as 3 weeks old. There were no significant differences between *Nlrp5 +/-* and *Nlrp5 +/+* oocytes (Figure 2B, C).

**Figure 2:**
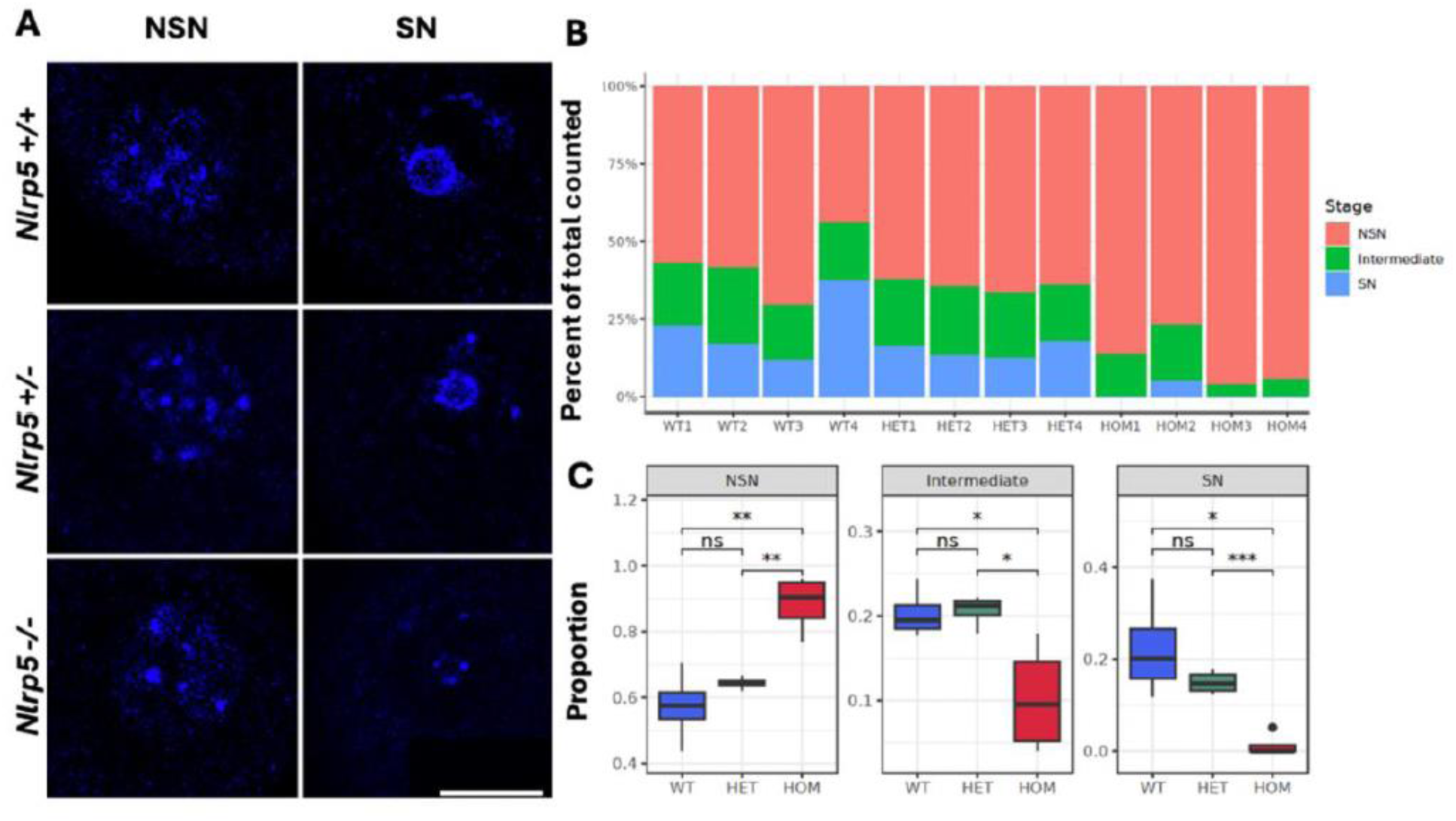
Analysis of nuclear maturation in *Nlrp5 -/-* oocytes. **(A)** Representative images of Hoechst-stained NSN and SN *Nlrp5 -/-, +/-* and *+/+* oocytes. Scale bar 20µm. **(B)** Proportions of GV oocytes of each stage per biological replicate (4 mice per genotype, 17-89 GV oocytes classified per replicate). **(C)** Proportion of GV oocytes of each stage per genotype group. T-test significance * p= <0.05, ** p= <0.01, *** p= <0.001. WT = wild-type, HOM = *Nlrp5 -/-*, HET = *Nlrp5 +/-*.

To detect differences in gene expression that occur between comparably staged GV oocytes, we collected GV oocytes from 3-week-old females of each genotype, and performed single-cell RNA sequencing. Because of the observed impairment of nuclear maturation in *Nlrp5-/-* oocytes, NSN/SN staging was inferred bioinformatically using a list of NSN-SN stage-specific DEGs^5^ (Figure S1(i), Table S1(i)). After filtering samples for minimum read count, between 2 and 6 samples per stage and genotype remained for analysis (Figure S1(ii)A). There was no difference in the mean number of genes detected in *Nlrp5 -/-* or *Nlrp5 +/-* oocytes at either stage (Figure S1(ii)B)). Global transcriptomic analyses demonstrated that *Nlrp5 -/-* oocytes could be separated from *Nlrp5 +/+* oocytes of the equivalent NSN/SN stage (Figure 3A), although when staging is not considered, this distinction is obscured.

**Figure 3:**
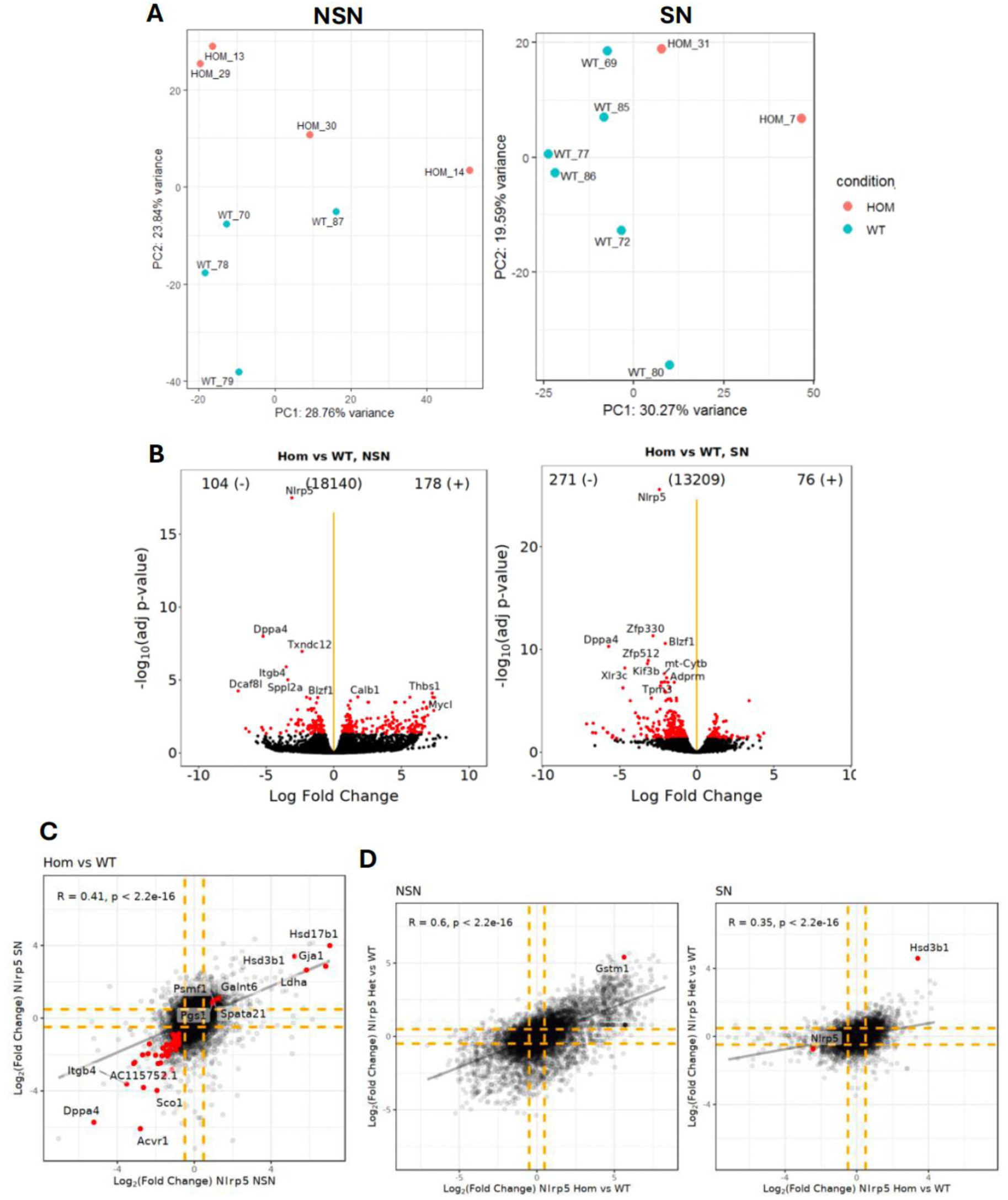
Comparison of *Nlrp5* -/- and *Nlrp5 +/+* oocyte transcriptomes, accounting for NSN/SN staging. **(A)** PCAs of *Nlrp5 +/+* (WT) and *Nlrp5 -/-* (HOM) samples (all expressed genes), after assignment to NSN (left) and SN (right) stages. **(B)** Volcano plots showing *Nlrp5 -/-* DEGs at NSN (left) and SN stage (right) respectively. Significant DEGs highlighted in red (significant where log2 FC is <-0.5 or >0.5, and BH-adjusted p value is <0.05), **(C)** Scatter plot showing correlation of Log2 FC gene expression for all genes detected in *Nlrp5 -/-* samples, comparing expression at NSN and SN stages. DEGs significant in both stages are highlighted in red (45 shared DEGs). FDR <= 0.05, lof2FC >=0.5 in either or both NSN and SN groups, R (correlation coefficient)=0.41. **(D)** Scatter plots showing positive correlation of Log2 FC gene expression in *Nlrp5 -/-* and *Nlrp5 +/-* samples, compared with *Nlrp5 +/+* samples. NSN and SN stages plotted separately. R (correlation coefficient)=0.6 and 0.35 respectively. DEGs significant in both *Nlrp5 +/-* and *Nlrp5 -/-* cohorts highlighted in red.

Differentially expressed genes (DEGs) detected between *Nlrp5 -/-* and *Nlrp5 +/+* oocytes (log2 FC <-0.5 or >0.5, Benjamini-Hochberg-adjusted p value <0.05) accounted for <3% of expressed genes: 282/18140 (1.55%) in NSN and 347/13209 (2.63%) in SN oocytes (Figure 3B, Table S1(ii)). DEGs were distributed across the whole range of transcript abundances (Figure S1(iii)). Taken together, this suggests that disruption of the SCMC in the absence of NALP5 is unlikely to lead to major effects on mRNA stability in GV oocytes, though some transcript specific effects on stability could explain a higher number of DEGs being downregulated in the more mature, transcriptionally inert, SN-stage oocytes than NSN oocytes (Figure 3B).

Although only 45 DEGs were called in common between NSN and SN stages, differential expression of DEGs at either stage was mostly correlated between stages (Pearson correlation, R=0.41, p<2.^2e-1^^6^; Figure 3C).

Far fewer DEGs were detected between the *Nlrp5 +/-* and *+/+* oocytes using the cut- offs above (Table S1(iii)), although fold-change in transcript abundances in the heterozygotes was positively correlated with fold-change in the homozygotes (Pearson correlation, R=0.6, p<2.^2e-16^ in NSN, R=0.35, p<2.^2e-16^ in SN; Figure 3D, FigureS1(iv)A, B). This indicates that a certain level of transcriptome misregulation is detectable in the *Nlrp5 +/-* oocytes, but given that *Nlrp5 +/-* oocytes can be successfully fertilised to produce viable embryos, the transcriptome misregulation detected does not reflect impaired developmental competence of heterozygotes oocytes.

Gene enrichment analysis highlighted general terms such as ‘cellular response to stimulus’, ‘steroid biosynthesis’ and ‘cellular anatomical entity’ in upregulated NSN-stage *Nlrp5 -/-* DEGs (Figure S1(v)A). The ‘Hsp110-Hsc70-Hsp25’ heat shock protein complex was enriched in upregulated DEGs at the SN stage, in addition to ‘steroid biosynthesis’ (Figure S1(v)B). Terms related to DNA binding and transcriptional regulation were enriched in downregulated DEGs at the SN stage (Figure S1(v)C). Apart from strong down-regulation of *Nlrp5* (Figure 3B), which might indicate nonsense-mediated decay of the mutant transcript, there was no evidence for altered mRNA abundance of components of the SCMC. And, other than *Dppa4*, no transcripts coding for known epigenetic modifiers were significantly differentially abundant (Table S1(ii)).

### Proteomic analysis shows misregulation of a subset of SCMC proteins in *Nlrp5 -/-* GV oocytes

Mass spectrometry analysis was performed on *Nlrp5 -/-, +/-* and *+/+* GV oocytes to determine if causes of the developmental defect could be detected in the proteome. GV oocytes were collected from 3-week-old mice in bulk samples (33 oocytes per sample) and the gel-LC-MS/MS analysis used data-independent MS/MS mode. The data were searched against the Uniprot canonical mouse proteome database using DIA-NN software. Over 5228 proteins were identified per sample (Figure S2(i), Table S2(i)). *Nlrp5 -/-* bulk oocyte samples showed a log_2_ fold change for NALP5 of -8.47 compared with *Nlrp5 +/+* oocytes, which corresponds to 0.3% of the *Nlrp5 +/+* level.

*Nlrp5 +/-* samples had a far lower but still significant log_2_ fold change of -0.55, corresponding to 68% of the *Nlrp5 +/+* NALP5 level. *Nlrp5 -/-* samples separate out from *Nlrp5 +/+* samples based on global protein abundances, with *Nlrp5 +/-* samples overlapping with both groups (Figure S2(i), Table S2(i)). To test whether the effects seen in the *Nlrp5 -/-* proteomics data could be attributed to a staging skew, the data were compared to existing wild-type SN/NSN proteomics data^48^; however, no significant difference in abundance in proteins that were known to be differentially abundant in NSN compared to SN was found (Figure S2(ii)).

When *Nlrp5 -/-, Nlrp5 +/-* and *Nlrp5 +/+* samples were clustered based on abundances of SCMC proteins, *Nlrp5 -/-* samples separate out strikingly from *Nlrp5 +/-* and *Nlrp5 +/+* samples (Figure 4A), although similar trends in SCMC protein abundances apply in the *Nlrp5 +/-* samples, but to a lesser degree (Table S2(i)). Significance cut-offs of log_2_ fold change <-0.5 or > 0.5 and false discovery rate (FDR) < 0.1 were used to determine significance of protein abundance differences between *Nlrp5 -/-* and *Nlrp5 +/+* oocyte samples. The -8.47 log_2_-fold reduction of NALP5 in *Nlrp5 -/-* samples is associated with a significant reduction in the abundances of seven known or putative components of the SCMC: TLE6, OOEP, ZBED3, KHDC3, NLRP4f, NLRP4b and NLRP9a. Of these, NLRP4b and NLRP9a have not previously been demonstrated to be part of the SCMC. The strong reduction (>85%) in TLE6 and OOEP, two proteins considered necessary for the complex to form^69^, at the protein level but not at the transcriptome level, demonstrates that the integrity of the complex is strongly affected by the ablation of NALP5. Notably, PADI6, NLRP2 (Q4PLS0), NLRP14, and five other NLRPs detected were not significantly changed in abundance in the *Nlrp5 -/-* samples (Figure 4A).

**Figure 4:**
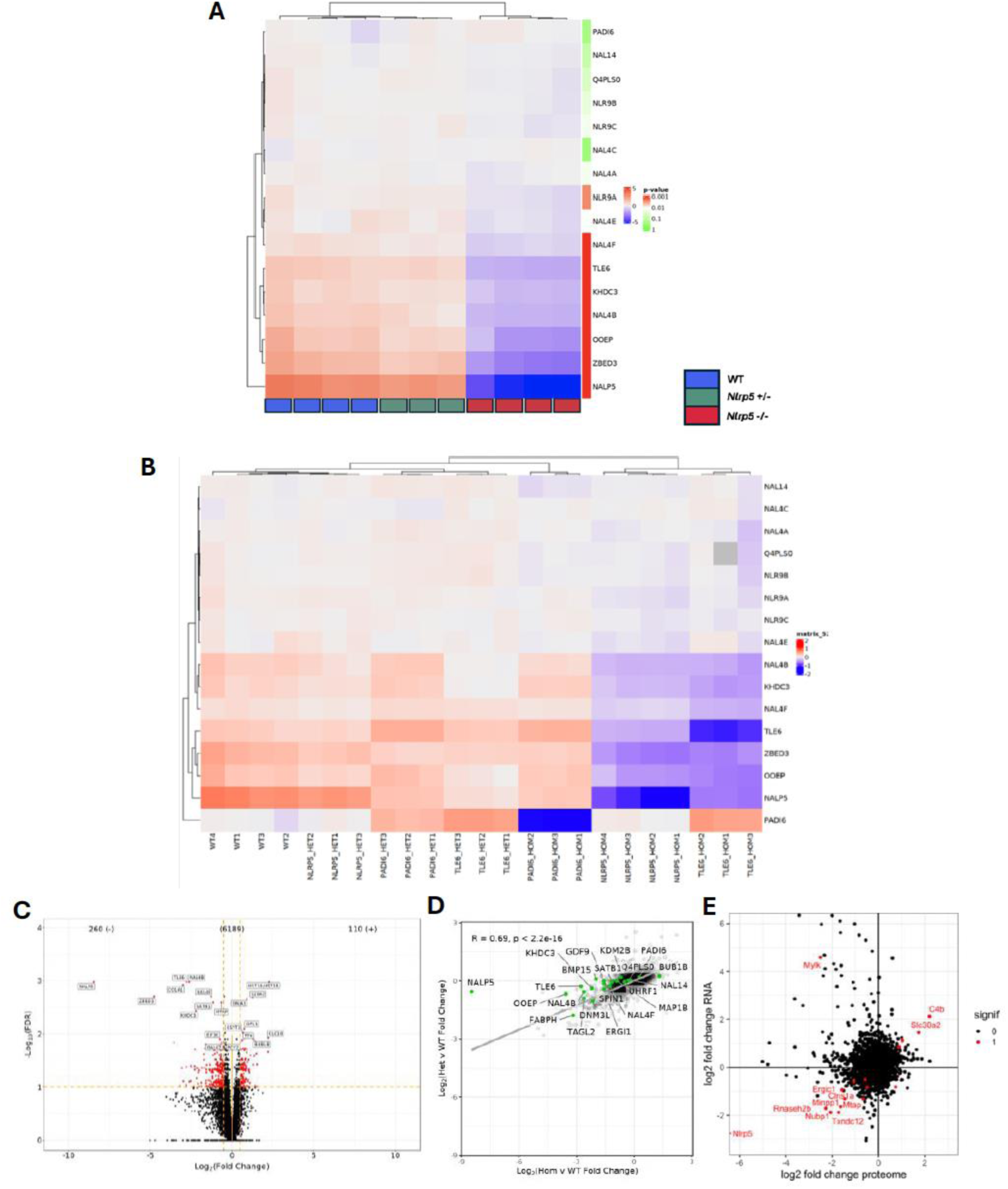
Comparative analysis of *Nlrp5-, Tle6-,* and *Padi6*-null GV oocyte proteomes. **(A)** Hierarchical clustering of raw protein abundances between *Nlrp5-/-* (RED), *Nlrp5 +/-* (GREEN), and *Nlrp5 +/+* (BLUE) samples. Heatmap showing Log2 abundance values with mean subtracted, for known and putative SCMC proteins. **(B)** Heatmap showing SCMC protein abundances across *Nlrp5-, Tle6-*, and *Padi6*-null datasets. Data were log-normalised, adjusted by subtracting the mean and dividing by the variance across all proteins for each dataset, and further normalised by subtracting the row means for each dataset. **(C)** Volcano plot of all differentially abundant proteins between *Nlrp5 -/-* and *Nlrp5 +/+* GV oocytes. Significant proteins highlighted in red, top 20 labelled (FDR < 0.1, log2 fold change of <-0.5 or > 0.5 [protein abundance altered by 30% or more]). **(D)** Scatterplot showing correlation of global protein abundances between the *Nlrp5 -/-* vs WT and *Nlrp5 +/-* vs WT comparisons (Log_2_ fold changes). Proteins of interest highlighted in green. **(E)** Scatterplot of global protein vs transcriptomic changes (log2 FC). Significant hits in both datasets highlighted in red.

Comparison of our *Nlrp5* proteomic data with published *Tle6-* and *Padi6*-null datasets^3^ shows greater similarities between the *Nlrp5 -/-* and *Tle6 -/-* proteomes than comparisons of either with the *Padi6* -/- proteome, when comparing log-normalised abundances of known and putative SCMC proteins (Figure 4B). This suggests two functionally separate (or at least partially redundant) SCMC protein subgroups. The often subtle but nonetheless measurable, change in abundance of the array of NLRP4 and NLRP9 isoforms across studied SCMC mutant proteomes indicates that these proteins are SCMC-related, although do not fit neatly into either of the two clear classes. Despite operating as separate functional subgroups to influence abundance of other SCMC-related proteins and other maternal proteins, both the NALP5-TLE6 category of SCMC proteins and the PADI6 category are thought to be necessary for the integrity of CPLs in the oocyte cytoplasm^32,34,42,49^, and null mutations in any of these three genes have similar 2-cell arrest phenotypes^34,43,45^.

### Broad range of maternal proteins altered in abundance in *Nlrp5*-null oocytes

370 differentially abundant proteins (DAPs) were identified in the *Nlrp5*-null data between *Nlrp5 -/-* and *Nlrp5 +/+* samples (Figure 4C). However, when the same significance cut-offs were used (<0.1 FDR, log_2_ fold change <= -0.5 or >= 0.5) in the *Tle6*- and *Padi6*-null datasets, only 10 DAPs were identified in the *Tle6*-null data, and 128 in the *Padi6*-null data. Further comparative analysis of the *Nlrp5*-null, and *Tle6*- and *Padi6*-null proteomes was performed using these three lists of DAPs. A published list of DAPs from an *Nlrp14*-null oocyte dataset^36^ was also used for comparison where overlapping proteins were detected.

Of the differentially abundant proteins (DAPs) between *Nlrp5 -/-* and *Nlrp5 +/+* samples, 260 were significantly reduced in abundance in *Nlrp5 -/-* samples and 110 were significantly increased (Figure 4C). Comparison of *Nlrp5 +/-* and *Nlrp5 +/+* samples yielded no significant DAPs using the same cut-offs, however, correlation analysis performed on the log_2_ fold change of all detected proteins concluded that, while not reaching the threshold for significance and with a smaller magnitude, the log_2_ fold changes were positively correlated between the homozygous and heterozygous *Nlrp5* mutant oocytes (Pearson correlation, R=0.69, p<2.2e^-16^; Figure 4D). Importantly, although *Nlrp5 -/-* and *Nlrp5 +/+* samples can be clearly separated based on both the transcriptome and proteome, with very few exceptions, no correlation exists between transcript and protein abundance changes (Figure 4E). Enrichment analysis for DAPs in the *Nlrp5 -/-* vs *Nlrp5 +/+* comparison resulted in very few hits. Upregulated DAPs were enriched for ‘regulation of protein modification process’, and downregulated DAPs were enriched for ‘Khdc3-Nlrp5-Ooep-Tle6 complex’ (Figure S2(iii)). As very few enrichment terms emerged from the enrichment analysis of the *Nlrp5 -/-* DAPs, published literature describing proteins and biological processes altered in SCMC mutant oocytes, coupled with hits from comparative analyses with other SCMC mutant datasets^3,36^, were used as guidelines to investigate changes in certain protein groups within the *Nlrp5*-null dataset.

DAPs in the *Nlrp5*-null oocytes included a broad range of proteins important for a diversity of biological processes necessary for oocyte and embryo development (Table S2(ii)). SCMC components have been linked to maintenance of euploidy during cleavage-stage mouse embryogenesis via regulation of microtubule-related proteins such as γ-tubulin. Significant reductions in abundances of tubulins and tubulin-associated proteins, and a reduction in acetylation of tubulins, have been described in *Padi6-* and *Nlrp14*-null oocytes ^3,36,49^. Three cytoskeleton-associated proteins were significantly reduced in *Nlrp5 -/-* samples: TBAL3, which was reduced by ∼30%; TAGL2, reduced by 80%, and MAP1B, reduced by over 40%. The only overlapping protein between the datasets was MAP1B, which was altered in both *Nlrp5-*null and *Padi6*-null oocytes. It was also listed as significantly reduced in *Nlrp14*-nulls^36^. Its abundance was unchanged in *Tle6*-nulls. Despite minimal overlap in effect on abundances of specific cytoskeleton-related proteins between *Nlrp5, Tle6*, and *Padi6*-null datasets, there is likely to be some overlapping function in cytoskeleton architecture, particularly via microtubule organisation.

Two proteins with important roles in folliculogenesis, GDF9 and BMP15, were strikingly reduced in *Nlrp5 -/-* oocytes (65% and 75%, respectively). GDF9 and BMP15 are thought to cooperate synergistically to influence granulosa cell proliferation and ovulation rate^50^, so an effect on one may influence the abundance of the other. *Gdf9*-null mice undergo arrest of folliculogenesis at the primary stage^51^. Interestingly, despite the strong reduction in GDF9 abundance in *Nlrp5 -/-* oocytes, no apparent difference in ovarian or follicular morphology, or ovulation rate in response to stimulation with exogenous gonadotrophins is observed in *Nlrp5* -/- mice^45^. *Bmp15-*null mice do not show evidence of reduced folliculogenesis, but are subfertile due to defective ovulation and reduced viability of embryos^52^. Tightly controlled *Gdf9:Bmp15* mRNA ratio and post-translational modifications of their protein products is considered important for controlling ovulation rate in a species-specific manner^53^. Although infertile, *Nlrp5*-null female mice have normal ovulation^45^. Despite these effects in *Nlrp5 -/-* oocytes, none of the other SCMC knockout datasets (*Tle6-, Padi6-, Nlrp14*-nulls) showed any significant change in GDF9 or BMP15 abundance.

Several factors involved in meiotic control and chromatin organisation were altered in *Nlrp5 -/-* oocytes, including SPIN1, SATB1 and BUB1B. Of these, SPIN1 and SATB1 were downregulated, and BUB1B was strongly up-regulated. SPIN1 is a chromatin reader^54^ and regulator of oocyte meiotic resumption, which may act by regulating maternal transcripts^55^. It was reduced by over 65% in *Nlrp5 -/-* samples, and by over 40% in *Tle6 -/-* samples, but was unaltered in *Padi6 -/-* samples. It was not listed as significantly altered in *Nlrp14*-null oocytes. SATB1 and BUB1B were not significantly changed in *Tle6*- or *Padi6*-null datasets, and were not listed in the *Nlrp14*-null DAP list.

Other significantly altered proteins in *Nlrp5*-null oocytes included FABP3, which binds free long-chain fatty acids (LCFAs) and transports them for cell metabolism, thereby protecting against lipid toxicity^56^, and ERGIC1, which is involved in protein cycling and is thought to play a role in protein transport between the endoplasmic reticulum and Golgi^57^. While most DAPs in the *Nlrp5 -/-* oocytes were downregulated, notable exceptions were heat-shock chaperone proteins HS90A, HS105, HS71A and HS71B (65%-380% increase), and apoptosis-related factors, such as TP63. The latter two groups may reflect impairments in cellular regulation and reduced oocyte developmental competence. However, none of the same heat shock family proteins were differentially abundant in *Tle6-* or *Padi6-*null samples. These differences may indicate cellular and developmental defects unique to *Nlrp5 -/-* oocytes.

### Epigenetic modifier proteins amongst those significantly altered in abundance in *Nlrp5*-null oocytes

When samples were plotted based on raw abundances of 11 epigenetic modifier proteins with known roles in *de novo* methylation and methylation maintenance, *Nlrp5 -/-* samples cluster separately from *Nlrp5 +/-* and *Nlrp5 +/+* samples (Figure 5A). While most of these proteins were not significantly altered in abundance in *Nlrp5 -/-* samples, three were: DNMT3L (>75% reduction compared with *Nlrp5 +/+*), UHRF1 (30% reduction) and KDM2B (>40% reduction) (Figure 5A, Table S2(i)). These reductions occurred without parallel changes in transcript level (Table S1(ii)).

**Figure 5:**
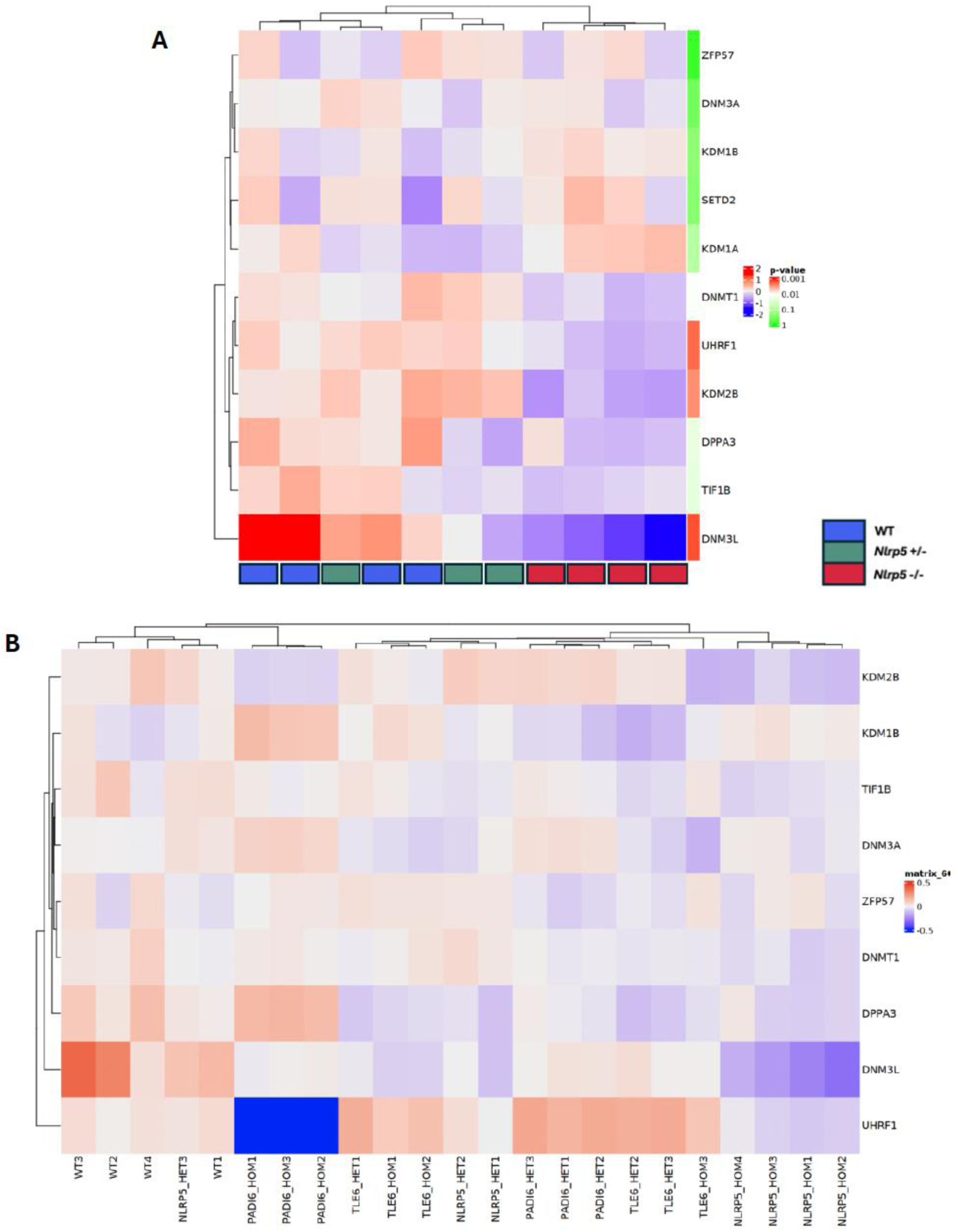
Altered abundances of known epigenetic modifier proteins detected in *Nlrp5* mass spectrometry data. **(A)** Heatmap of Log2 abundances with mean subtracted. RED = *Nlrp5 -/-,* GREEN = *+/-,* BLUE = *+/+.* **(B)** Heatmap showing abundances of known epigenetic modifier proteins across *Nlrp5, Tle6*, and *Padi6* datasets. Data were log-normalised, adjusted by subtracting the mean and dividing by the variance across all proteins for each dataset, and further normalised by subtracting the row means for each dataset.

DNMT1 also appeared reduced in *Nlrp5 -/-* samples, but did not meet the cut-off for significance. These key proteins also show some changes in other datasets. Thus, DNMT3L, an important cofactor for DNMT3A for *de novo* DNA methylation activity in the mouse, is reduced by 30% in *Padi6 -/-* oocytes^3^, but not in *Tle6 -/-* oocytes (Table S2(i)). UHRF1 is a factor important for maintenance of DNA methylation via the localisation of DNMT1 to hemi-methylated sites of the DNA during the S phase of DNA replication^15^. Its abundance was not significantly affected in *Tle6*-null GV oocytes, but it is strongly reduced in *Padi6-*null (Figure 5B; Table S2 (i)), and *Nlrp14*- null oocytes^36^. KDM2B, a histone H3 demethylase known to target H3K36me2, was significantly reduced in *Nlrp5*-null as well as in *Padi6*-null oocytes, but unchanged in *Tle6-*nulls (Figure 5A, B).

### DNMT3L and UHRF1 localisation altered in *Nlrp5* KOs, while DNMT1 localisation unchanged

Due to the finding that DNMT3L is reduced in abundance by over 75% in *Nlrp5 -/-* oocytes, we tested whether DNMT3L localisation is altered, by immunofluorescence. Because of the low numbers of *Nlrp5 -/-* SN oocytes, all cross-genotype comparisons were based on staining of NSN oocytes. Relative mean nuclear fluorescence and relative mean cytoplasmic fluorescence of DNMT3L were not significantly different between genotypes (Figure S2(iv)). However, when the nuclear-to-cytoplasmic DNMT3L fluorescence ratio was calculated for each oocyte, the ratio was significantly altered in *Nlrp5 -/-* oocytes compared to *Nlrp5 +/+* and *Nlrp5 +/-* oocytes (Figure 6A, B), with relatively less DNMT3L fluorescence in the nucleus.

**Figure 6:**
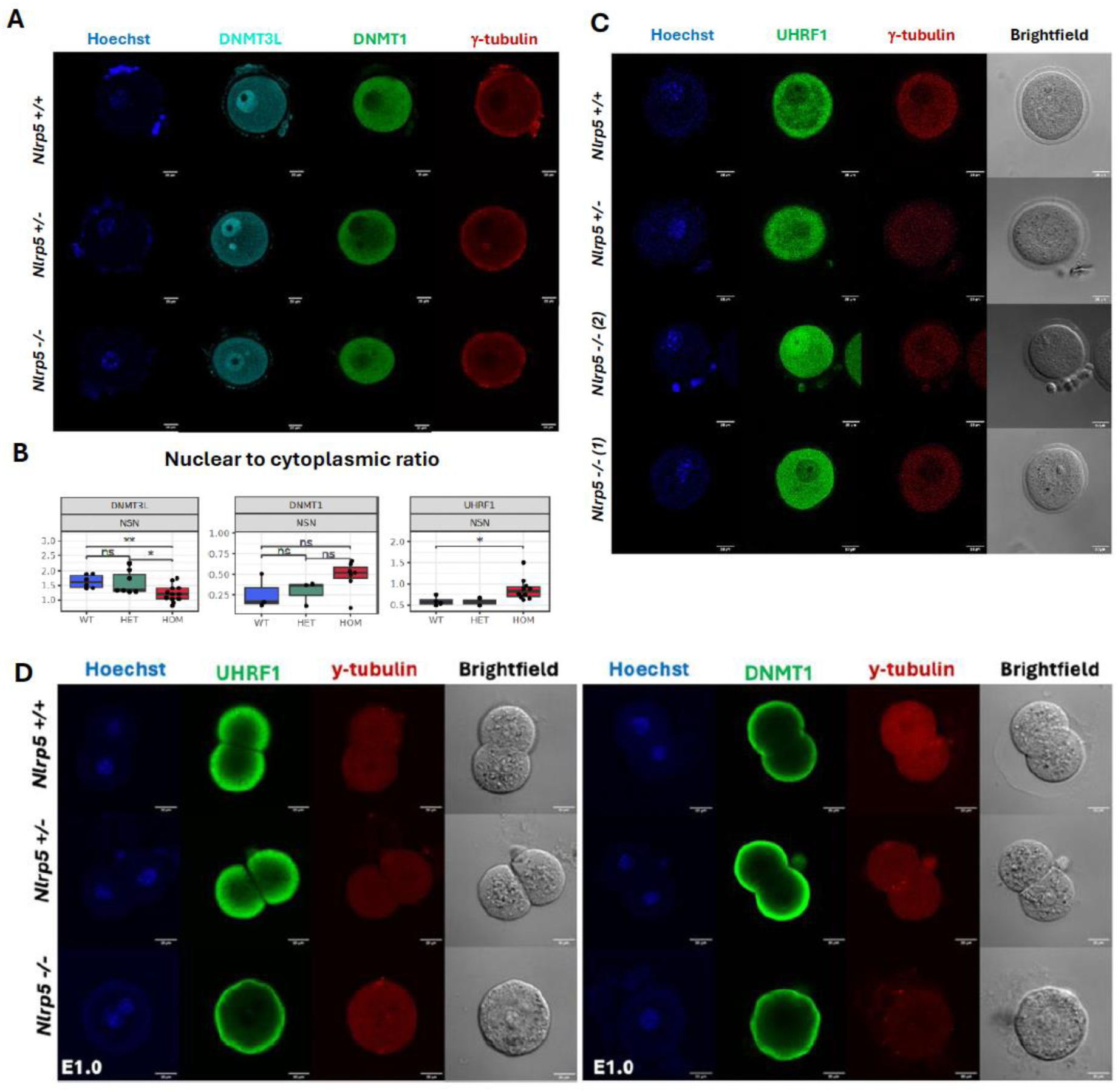
Immunofluorescence analysis of DNMT3L, DNMT1 and UHRF1 in *Nlrp5 -/-* GV oocytes and E1.0 maternal *Nlrp5 -/-* embryos. **(A)** Representative immunofluorescence images of NSN-stage GV *Nlrp5 -/-, +/-* and *+/+*. Oocytes with DNMT3L (cyan), DNMT1 (green). **(B)** Box-whisker plots showing nuclear to cytoplasmic fluorescence ratio for DNMT3L (*n = 12 Nlrp5 -/-, 7 Nlrp5 +/-, 6 Nlrp5 +/+),* DNMT1(n = 7 *Nlrp5 -/-, 3 Nlrp5 +/-, 3 Nlrp5 +/+*), and UHRF1 (n = 10 *Nlrp5 -/-,* 2 *Nlrp5 +/-, 4 Nlrp5 +/+*) for each genotype. Shapiro-Wilk normality test applied. If p>0.05, students t-test was used. If p<0.05, Wilcoxon Rank-Sum test was used. ** = significant (p-value <0.01), * = significant (p-value <0.05), ns = difference not significant. 20µm scale bar. **(C)** Representative immunofluorescence images of NSN-stage GV *Nlrp5 -/-, +/-* and *+/+*. Oocytes with UHRF1 staining (green). γ- Tubulin staining in red. All oocytes collected at 8 weeks old. **(D)** Representative immunofluorescence images of NALP5 maternal -/-, +/- and +/+ embryos, at E1.0. Blue = Hoechst DNA staining, green = UHRF1 (left), DNMT1 (right), red = γ- Tubulin. 20µm scale bar.

We also evaluated the localisation of DNMT1 and UHRF1, because these proteins have been reported to be mislocalised from the cytoplasm to the nucleus in *Padi6*- null GV oocytes and maternal *Padi6*-KO pre-implantation embryos, while DNMT1 is mislocalised to the nucleus in maternal *Nlrp14*-KO embryos^35,58^. There was no significant mislocalisation of DNMT1 evident in *Nlrp5 -/-* GV oocytes, with DNMT1 staining localised to the cytoplasm in oocytes of all genotypes (Figure 6A, B, Figure S2(iv)). UHRF1 was predominantly cytoplasmic in *Nlrp5 +/-* and *Nlrp5 +/+* oocytes although some was present in the nucleus. In *Nlrp5 -/-* oocytes, however, there was significantly increased nuclear UHRF1 staining (Figure 6B, C, Figure S2(iv)). When assessing embryonic day (E) 1.0 embryos generated by IVF from *Nlrp5 -/-* oocytes, we found that neither DNMT1 nor UHRF1 were altered in localisation compared with wild-type E1.0 embryos (Figure 6D). Note, however, smaller differences may be obscured by differences in embryo staging between the maternal *Nlrp5*-KO embryos and controls. This misregulation of epigenetic modifier proteins in mouse SCMC- mutants, including in *Nlrp5*, provides a biological mechanism for the methylation and imprinting phenotypes seen *in vivo*.

### Low level of global DNA hypomethylation in *Nlrp5 -/-* GV oocytes coinciding with hypomethylation over a subset of gDMRs

To understand the epigenetic effects of absence of NALP5, GV-stage oocytes from *Nlrp5 -/-* and *Nlrp5 +/+* females were processed via single-cell post-bisulphite adapter tagging (scPBAT-seq) to generate libraries for whole genome DNA methylation analysis^59,60^. Because DNMT3L is essential for *de novo* methylation in mouse oocytes^12,61^, the significant reduction of DNMT3L in *Nlrp5* -/- oocytes identified by mass-spectrometry suggested the possibility of impaired methylation. After filtering out samples with low read counts or suspected somatic-cell DNA contamination, 10 *Nlrp5 -/-* oocytes and 19 *Nlrp5 +/+* oocytes remained for analysis (Table S3(i)). Methylation was quantitated across 100-CpG consecutive tiles in the genome. The average global CpG methylation percentage was significantly lower (p- value <0.001) in the *Nlrp5* -/- compared to *Nlrp5 +/+* oocytes, with a difference of 4.98% between median global CpG methylation in *Nlrp5 -/-* and *Nlrp5 +/+* samples (Figure 7A). For further analysis, samples were pseudobulked for higher coverage of gene features. The *Nlrp5 -/-* pseudobulked cohort had a combined total of 31.6 million reads, and the *Nlrp5 +/+* pseudobulked cohort had a total of 28.2 million reads.

**Figure 7:**
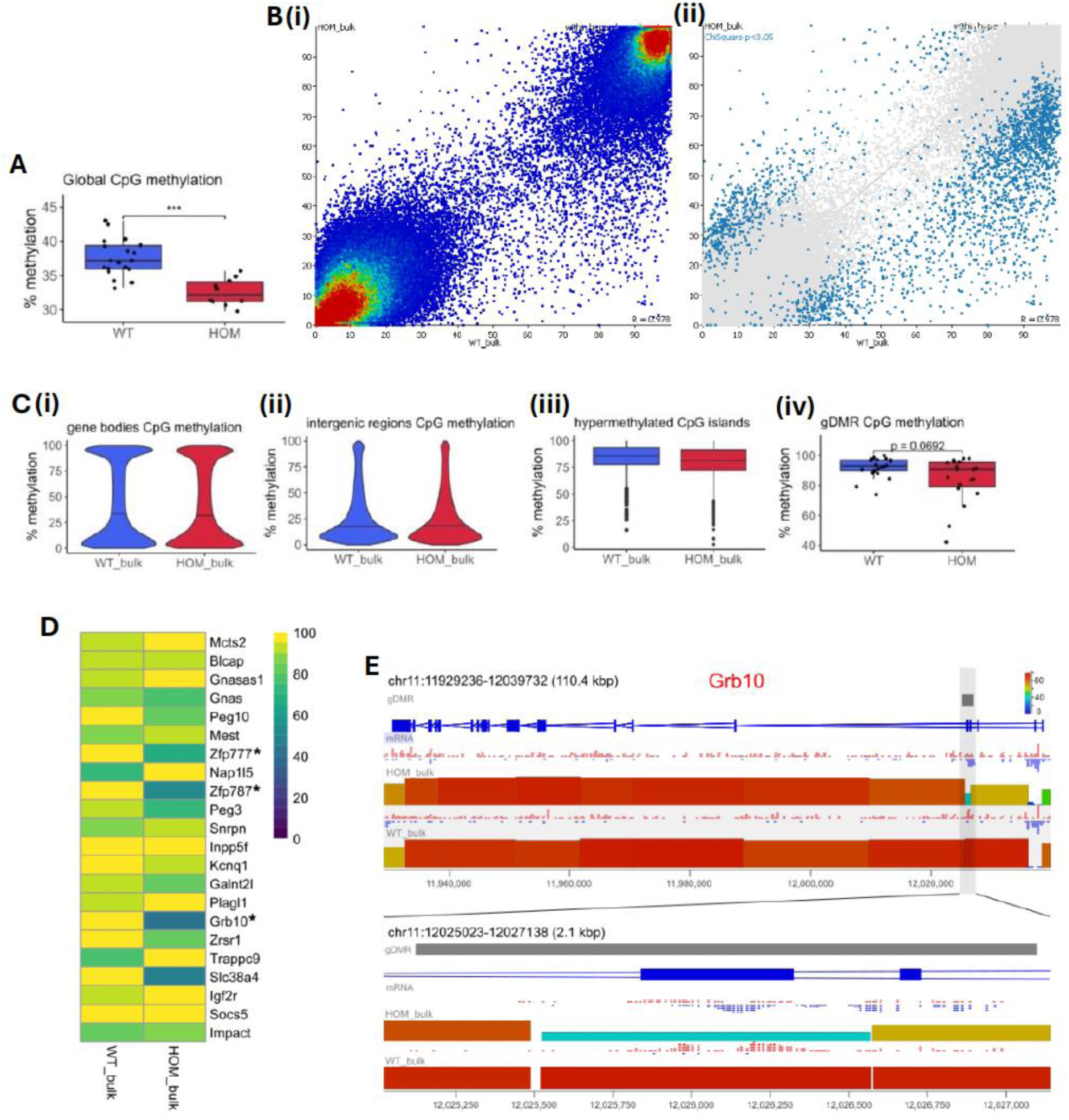
Global attenuation of DNA methylation overlapping with gDMRs. **(A)** Box-whisker plot showing global % CpG methylation of each WT (*Nlrp5 +/+)* and HOM (*Nlrp5 -/-)* oocyte. Welch t-test, *** = p value of <0.001 **(B)** (i) Scatterplot of bulked WT and HOM samples– all 100-CpG tiles, coloured for density. (ii) Scatterplot of tiles called differentially methylated (tiles with chi square p-value of <0.05). **(C)** Violin and box-whisker plots showing % methylation over certain genomic features: (i) Gene body CpG methylation, (ii) intergenic region CpG methylation, (iii) annotated hypermethylated CpG islands, and (iv) gDMR CpG methylation, Wilcoxon rank-sum test showing borderline but not significant p value (significance cutoff <= 0.05). **(D)** Heatmap showing % methylation at listed oocyte gDMRs in *Nlrp5 -/-* (HOM) and *Nlrp5 +/+* (WT) pseudobulked samples. * indicates significant under above cutoffs. **(E)** Example of hypomethylated gDMR: Seqmonk genome viewer view of the *Grb10* locus in *Nlrp5 -/-* and *Nlrp5 +/+* datasets, with (below) zoom-in on the *Grb10* gDMR. Each coloured block represents a 100-CpG tile, with height and colour-coding indicating % methylation of CpGs called across the tile. Methylation calls of individual CpG sites indicated above as red (methylated) or blue (unmethylated) ticks.

*Nlrp5 -/-* and *Nlrp5 +/+* samples recapitulated the typical bimodal methylation patterning unique to oocytes^8^ (Figure 7B), and percentage methylation at gene bodies, intergenic regions, and annotated hypermethylated CpG islands (CGIs) was not significantly different between *Nlrp5 -/-* and *Nlrp5 +/+* oocytes (Figure 7C). To test for sites of differential methylation, a chi-square test was performed on 100-CpG-tiles within annotated oocyte hyper- and hypomethylated domains^62^ (significance < 0.05 after multiple testing correction, minimum percentage difference = 25%). The test produced 2452 differentially methylated hits between the *Nlrp5* -/- and *Nlrp5 +/+* groups (Figure 7B(ii)). When assessing methylation at all annotated oocyte gDMRs, there was no significant difference between genotypes across all gDMRs (Figure 7C(iv)). However, 3 individual gDMRs (*Zfp777*, *Zfp787* and *Grb10;* Figure 7D, E) were significantly hypomethylated in the *Nlrp5 -/-* group compared to *Nlrp5 +/+* oocyte probes, as determined by chi-square test on 100-CpG probes overlapping with gDMRs (Table S3(ii)).

To confirm that the observed global hypomethylation in the *Nlrp5* -/- oocytes could not be attributed to a skew in the NSN-SN staging ratio, a comparative analysis was performed on a wild-type NSN/SN mouse GV oocyte staging methylation dataset^63^. The staging data were quantitated using the same 100-CpG windows across the genome. When principal component analysis was performed on samples from the *Nlrp5* and staging datasets, *Nlrp5 -/-* samples separated out from all wild-type samples, including those of the SN/NSN staging dataset. Although there was some effect of staging, there was limited overlap of *Nlrp5 -/-* samples with NSN samples (Figure S3). Further, none of the 3 gDMRs identified as differentially methylated in *Nlrp5* -/- GV oocytes were significantly differentially methylated in the staging dataset.

## Discussion

Multi-omic analysis of GV oocytes from our CRISPR/Cas9-generated *Nlrp5 -/-* mice demonstrates that SCMC integrity is lost, making *Nlrp5* mutants an ideal model in which to study overall SCMC function. We find that in *Nlrp5* -/- GV oocytes, the abundance and/or localisation of some key epigenetic factors is altered, and there is a modest global hypomethylation, which encompasses some gDMRs. Additionally, the ratio of NSN-SN stage oocytes in the *Nlrp5 -/-* cohort skews towards the less mature NSN-stage, detectable as early as 3 weeks old. While this staging skew likely contributes to some of the methylation differences observed, we show that it does not account for all of the differences in DNA methylation. Indeed, the three significantly hypomethylated gDMRs detected in *Nlrp5 -/-* GV oocytes are not significantly hypomethylated in wild-type NSN GV oocytes. Immunofluorescence staining of GV oocytes showed that, while NALP5 was ablated in *Nlrp5 -/-* GV oocytes, it was present throughout the cytoplasm in *Nlrp5 +/-* and *Nlrp5 +/+* oocytes, not just at the subcortex, supporting recent studies describing the presence of SCMC proteins throughout the cytoplasm^3^. *Nlrp5 +/-* oocytes lacked subcortical saturation of NALP5 fluorescence. Given that *Nlrp5* +/- oocytes are developmentally competent, the previously described subcortical localisation of NALP5 (and potentially other SCMC proteins) in wild-type oocytes is unlikely to be biologically relevant.

### Two functional subcategories of SCMC proteins exist

A comparative analysis with *Tle6*-KO oocytes^3^ demonstrated the same trends in reduction of SCMC proteins as the *Nlrp5* KO, suggesting similar effects of these mutants on integrity of a subset of SCMC proteins. Three known SCMC proteins were not altered in abundance in the *Nlrp5 -/-* oocytes: PADI6, NLRP14 and NLRP2. These three SCMC proteins were also unaltered in the *Tle6* KO. Comparative analysis with *Padi6* KO oocytes^3^ demonstrated the opposite trends: reductions in NLRP14 and NLRP2, but little to no effect on other SCMC proteins. These trends indicate that two functionally separate subgroups of SCMC proteins exist, the NALP5-TLE6 group, and the PADI6-dependent group. This is further supported by the global comparative analysis of these three datasets (*Nlrp5-, Tle6-, Padi6*-nulls) which shows a greater overall similarity between *Nlrp5 -/-* and *Tle6* -/- oocyte proteomes than either share with *Padi6* -/- oocytes. Two new candidates for SCMC membership were identified in the *Nlrp5* KO, NLRP4b and NLRP9a, both of which were significantly reduced in *Nlrp5 -/-* oocytes. Various proteins of the reproductive clade of the NLRP family^64^ appear to interact with other SCMC proteins with a degree of redundancy, as their abundances differ greatly, even between *Nlrp5* and *Tle6* KOs.

Previous studies have suggested that the SCMC may regulate the stability and availability of maternal proteins, potentially facilitating their accessibility to nuclear translocation factors at key timepoints in development ^3,31,32,34^. By investigating the overlapping roles of the SCMC and CPLs, the studies have argued that this may not only ensure the timely and efficient localisation of epigenetic modifiers (and other regulatory factors) to the nucleus, but may also protect them against their premature degradation by the proteasome. Indeed, the SCMC may act as a structural scaffold for storage and stabilisation of maternal factors necessary for development, and the presence of organised CPLs may simply reflect a functional SCMC.

In the present study, a subset of SCMC proteins, as well as three epigenetic modifiers, DNMT3L, UHRF1 and KDM2B, and a broad range of other proteins important for oocyte development, were significantly altered in abundance, reflecting a strong effect of the knockout on the *Nlrp5* -/- oocyte proteome. In contrast, there were no effects on SCMC or listed epigenetic modifier genes observable in the *Nlrp5* -/- transcriptome, and the overall correlation of DAPs with DEGs was low. While a level of misregulation is observable in both the proteome and transcriptome, the proteome shows a larger effect, and more specific changes to oocyte maternally stored factors. This indicates that a greater level of misregulation occurs post-translationally, and that the eventual 2-cell arrest phenotype observed in embryos derived from *Nlrp5 -/-* oocytes is likely a result of deficiencies in maternally stored proteins, arising from ineffective storage or localisation at key developmental timepoints. The *Nlrp5* KO affects the abundance of a wide variety of different proteins, from building blocks of the cytoskeleton and modifiers of heterochromatin to factors involved in protein transport between the ER and the Golgi, in nuclear transport, and in epigenetic modification. This variety of regulatory targets suggests a broad role of NALP5 and the SCMC in storage and sequestration of maternal factors in the oocyte, binding a wide variety of proteins to protect them from proteasomal degradation, premature assembly into organelles or functional complexes, or premature translocation into the nucleus. This is especially important in the oocyte, where, after transcriptional arrest at the SN GV stage, further oocyte development and early embryo development is entirely mediated by the pre-existing pool of maternal factors.

The differences in effects on the proteome in *Nlrp5*-, *Tle6*-, and *Padi6*-null oocytes suggest that some groups of proteins are preferentially bound by certain SCMC subgroups, whereas others may be specifically regulated by individual SCMC proteins. Similarities in downregulated tubulin-related proteins in *Padi6* and *Nlrp14*- nulls suggests that this subgroup of the SCMC has a shared function in microtubule- regulation independent of NALP5 and TLE6, although NALP5, independently from TLE6, seems to regulate the cytoskeleton via TBAL3 and TAGL2. UHRF1 appears to be preferentially regulated by the PADI6-NLRP14 subgroup, with stronger reductions of UHRF1 abundance in these mutants than in *Nlrp5* knockouts, although this and another recent study show interactions between NALP5 and UHRF1^65^. Other DAPs identified in the *Nlrp5* -/- samples, such as FABP3, ERGIC1, SATB1, GDF9 and BMP15, were not significantly altered in abundance in the other SCMC KO datasets, indicating that they are preferentially regulated by NALP5. FABP3 is a key factor in lipid metabolism and protection against lipid toxicity^56^. ERGIC1 is a cycling membrane protein with roles in protein transport between the endoplasmic reticulum and Golgi^57^. SATB1 is a chromatin organiser and transcription factor which regulates several cellular processes such as differentiation, proliferation, and apoptosis^66^.

GDF9 and BMP15 are both important in folliculogenesis and the regulation of ovulation. These are just a few of the proteins significantly altered in abundance in *Nlrp5* -/- oocytes, illustrating the variety of proteins and pathways potentially regulated by the SCMC and NALP5 individually.

### Roles for NALP5 in DNA methylation and genomic imprinting

Three epigenetic modifier proteins, DNMT3L, UHRF1, and KDM2B, were significantly reduced in *Nlrp5* -/- oocytes; *Nlrp5 -/-* oocytes also had proportionally less DNMT3L in their nucleus than cytoplasm, and proportionately more UHRF1 in their nuclei, suggesting that NALP5 affects DNMT3L and UHRF1 localisation as well as abundance. *Dnmt3l*-null female mice are viable but sterile, with offspring lethality attributed to abnormal maternal imprinting^67^. A 75% reduction in oocyte DNMT3L could affect *de novo* methylation via its influence on the catalytic activity of DNMT3A^7,21^, and levels of global DNA methylation would be expected to be lower than in wild-type GV oocytes. While the mild 30% reduction in total UHRF1 abundance in *Nlrp5 -/-* oocytes would not suggest a biological effect on UHRF1-mediated *de novo* methylation, the mislocalisation of the UHRF1 to the nucleus in these oocytes means that an effect of UHRF1 on levels of DNA methylation in these oocytes cannot be ruled out. One hypothesis is that a nuclear reduction of DNMT3L could be partially supplemented by a nuclear increase in UHRF1, resulting in a more subtle hypomethylation phenotype. Indeed, *Nlrp5* -/- oocytes showed a low but significant level of global hypomethylation that could not be explained purely by potential staging differences between *Nlrp5 -/-* and *Nlrp5 +/+* samples. This modest global hypomethylation may be due to an attenuation or delay in complete *de novo* DNA methylation as a result of low DNMT3L availability. In addition to the global hypomethylation observed in *Nlrp5 -/-* oocytes, several significantly hypomethylated regions were identified, including three that coincided with known imprinted gDMRs or that shared a promoter with a nearby gDMR. The limited number of gDMRs affected would suggest that these are the DMRs most susceptible to deficiencies in the *de novo* methylation apparatus.

While UHRF1 and DNMT1 can contribute to *de novo* methylation in oocytes, they are particularly crucial for maintenance of DNA methylation. UHRF1 was the most strongly reduced protein in *Padi6* and *Nlrp14* nulls (∼90%) aside from PADI6 and NLRP14, respectively. Furthermore, UHRF1 and DNMT1 were mislocalised to the nucleus in MII *Padi6*-null oocytes and maternal *Padi6*-null embryos, coinciding with mild genomic hypermethylation in the oocytes and a more dramatic hypermethylation in the embryos, likely due to excessive maintenance of DNA methylation marks and failure of demethylation. Similarly, in *Nlrp14* -/- oocytes UHRF1 was mislocalised to the nucleus (DNMT1 localisation was not studied), although the methylation was not significantly altered. However, the subtlety of the hypermethylation in *Padi6*-null oocytes suggests that a similar phenotype in *Nlrp14*-/- oocytes is still plausible. While UHRF1 was also mildly reduced (∼30%) in *Nlrp5 -/-* GV oocytes, it is not known to be haploinsufficient and thus taken together with there being no significant effect on DNMT1 abundance or localisation, it is unlikely to affect methylation maintenance in the *Nlrp5*-null. Thus, NALP5 appears to preferentially regulate *de novo* methylation, while PADI6 seems to instead preferentially regulate methylation maintenance, indicating a balancing role of the two SCMC subgroups in maintaining normal DNA methylation patterning, via regulation and timely localisation of either *de novo* DNA methylation establishment proteins, or maintenance factors.

KDM2B is a H3 demethylase known to target H3K4me3 and H3K36me2. It is part of the variant PRC1 complex, recruiting Polycomb-1 proteins to unmethylated CpGs^68,69^. It was significantly reduced by over 40% in *Nlrp5* -/- oocytes and significantly reduced by ∼30% in *Padi6* -/- oocytes. The significant reduction of this protein in the *Nlrp5* KO may influence epigenetic reprogramming indirectly by causing a lower level of Polycomb-1 protein recruitment to unmethylated CpG regions. Interestingly however, an oocyte-specific *Kdm2b* knockout does not adversely affect embryonic development^70^ so the biological relevance of KDM2B in oocytes remains to be determined. Another histone demethylase, KDM1B, was significantly increased in abundance in the *Padi6-*null dataset, and listed as significantly increased in *Nlrp14 -/-* oocytes^36^, but was not altered in *Nlrp5* or *Tle6*- nulls. Oocytes from KDM1B-deficient females show loss of methylation at CGIs, including most imprinted gDMRs^71,72^, and embryos derived from *Kdm1b* -/- oocytes die before mid-gestation with imprinted gene deregulation^71^. The increased abundance of KDM1B exclusively in *Padi6-* and *Nlrp14*-nulls may indicate a separate role of the PADI6-NLRP14 SCMC subset in epigenetic regulation.

The epigenetic modifier proteins altered in abundance in *Nlrp5* -/- oocytes demonstrate that a mutation in one core SCMC gene can potentially mis-regulate *de novo* establishment of DNA methylation (via DNMT3L and potentially UHRF1) in the mouse oocyte, DNA methylation maintenance (via UHRF1) in the pre-implantation embryo, or histone modifications (via KDM2B) depending on the degree of altered protein abundance or mislocalisation. The downstream effects of SCMC mutations on DNA methylation in the embryo or later development are likely determined by the composition of the repertoire of proteins misregulated by a given SCMC mutation, to varying degrees of penetrance depending on severity of mutation and the redundancies between some SCMC components.

These findings have important implications for SCMC defects in humans, where severe global hypomethylation has been reported in oocytes deficient in the SCMC component KHDC3L^37^. The substantial deficiency in DNMT3L we observe in *Nlrp5* -/- mouse GV oocytes provides a plausible mechanism for the modest global hypomethylation, including at susceptible imprinted regions. However, the mouse model does not satisfactorily explain the human phenotype, as DNMT3L is not expressed in human oocytes^73^. Therefore, while this work supports an important link between the SCMC and DNA methylation patterning via the stability and localisation of epigenetic modifiers, the varied and species-specific effect of each individual SCMC-subunit^35^ means that specific human phenotypes will have to be further explored in models more similar to the human.

### Conclusion

This study points towards a mechanistic explanation for the link between SCMC integrity in the oocyte, and faithful DNA methylation patterning. Through multi-omic profiling of our *Nlrp5* KO, coupled with comparative proteomics analysis with other SCMC mutants, we consider how mutations in the SCMC can lead to a plethora of different imprinting and DNA methylation disorders, depending on the SCMC component in question, and on the penetrance of the SCMC mutation. We demonstrate that the NALP5 mutant phenotype is a consequence of misregulation of stability of a variety of oocyte maternal factors, with detrimental effects on several different biological processes crucial to oocyte developmental competence. One of the processes affected is *de novo* DNA methylation establishment, and a low level of global hypomethylation is observed in the *Nlrp5* -/- oocyte genome, overlapping with DMRs coinciding with three known imprinted genes. This is likely due to the adverse effects of the NALP5 KO on the abundance and localisation of DNMT3L, a key cofactor for murine *de novo* DNA methylation.

## STAR Methods

### RESOURCE AVAILABILITY

#### Lead Contact

Further information and requests for resources and reagents should be directed to Gavin Kelsey.

#### Data and Code Availability

Single-cell RNA and PBAT sequencing data have been deposited at Gene Expression Omnibus database and is available as of the date of publication under accession number: GSE278144.

Mass spectrometry (Orbitrap Eclipse (ThermoFisher Scientific, UK)) data have been deposited to the ProteomicXchange Consortium via the PRIDE partner repository with the dataset identifier PXD056859, available as of the date of publication.

This paper does not report original code.

Any additional information required to reanalyze the data reported in this paper is available from the lead contact upon request.

### EXPERIMENTAL MODEL AND SUBJECT DETAILS

All animal experimental procedures were approved by the Animal Welfare and Ethical Review Body at the Babraham Institute and were conducted under authority of the UK Home Office issued licences in accordance with the Animal (Scientific Procedures) Act 1986.

### METHOD DETAILS

#### Generation and verification of *Nlrp5* knockout mouse line

The *Nlrp5* knockout (KO) mouse model was generated using CRISPR/Cas9 technology by directly targeting mouse zygotes. Specifically, a single guide RNA (sgRNA) was designed using Benchling (https://www.benchling.com/crispr), and the guide sequence TGTCGAACTTAGCAATCACA was identified within a constitutive exon and essential protein domain. The CRISPR editing was performed by Asif Nakhuda of the Babraham Institute Gene Targeting Facility. Mouse zygotes were obtained from superovulated C57BL6/Babr females, and Cas9 protein (100 ng/µl, IDT) along with sgRNA (100 ng/µl, IDT) diluted in Opti-MEM was introduced into the zygotes via electroporation using the NEPA21 device (Sonidel, Japan) with the following settings: 40V, 3.5 ms pulse lengths, 50 ms intervals, and 4 pulses. The embryos were cultured overnight, and on the following day, 2-cell stage embryos were transferred into pseudopregnant CD1 females. Ear clips were collected from the F0 pups post-weaning for genotyping to identify KO genotypes. Genotyping PCR amplification was conducted over the Cas9/sgRNA-induced DSB using the primers Forward 5’-CCATTCAGGTTTCCTCCCAT-3’ and Reverse 5’- CCTGGTCTTCCATGTAGGAT-3’. Samples were purified using the Zymo DNA extraction kit (Zymo Research, USA) and sent for Sanger sequencing (Azenta, USA). Sanger sequencing traces were analysed using Inference of CRISPR Edits (ICE; Synthego, USA) to identify F0 mice with KO genotypes. These F0 mice were then crossed with wild-type C57BL6/Babr females to produce F1 offspring.

Genotyping was performed by Transnetyx (USA) using a TaqMan-based assay to obtain real-time PCR data for automated detection of the desired frameshift mutation. Subsequent litters were also genotyped via Transnetyx using the same TaqMan-based assay.

#### Timed matings and animal husbandry (litter counts/sizes)

All mice used in this study were bred and maintained in the Babraham Institute Biological Support Unit (BSU). Ambient temperature was ∼19-21°C and relative humidity 52%. Lighting was provided on a 12-hour light: 12-hour dark cycle including 15 min ‘dawn’ and ‘dusk’ periods of subdued lighting. After weaning, mice were transferred to individually ventilated cages with 1-5 mice per cage. Mice were fed CRM (P) VP diet (Special Diet Services, UK) *ad libitum* and received seeds (e.g. sunflower, millet) at the time of cage-cleaning as part of their environmental enrichment. Animal husbandry was managed by the BSU, including the setting up of timed matings, and routine genotyping of young mice, (via Transnetyx), once the *Nlrp5* knockout colony had been established.

#### Oocyte and embryo collections

Female mice were sacrificed at 3 weeks old, and ovaries were dissected out and collected in M2 medium (Sigma Aldrich, USA, cat:M7167) on ice. Sacrificed *Nlrp5 -/-* and *Nlrp5 +/-* females were always littermates, and *Nlrp5 +/+* females usually from the wild-type colony. For immunofluorescence and proteomics analysis, germinal vesicle-stage (GV) oocytes were isolated from ovaries in M2 medium using the scratch method, whereby oocytes were released from the ovary by gently scratching the ovary with a fine needle. Oocytes were manually washed in 4-5 drops of M2 medium using a drawn-out mouth pipette. Oocytes were then rinsed in PBS 3 times to minimise bovine serum albumen (BSA, from the M2 medium) contamination during mass spectrometry analysis. They were then either fixed in 4% paraformaldehyde (PFA; Merck, USA) for 15 mins for downstream immunostaining and stored in PBS-Tween at 4°C, or snap frozen in 5μl PBS for mass-spectrometry analysis. For single cell methylation and transcriptome sequencing, oocytes were instead collected using a collagenase digestion method (to reduce chance of somatic-cell contamination). The digestion mix was prepared as follows, for each pair of ovaries: 454.2μl DPBS (with Mg2+. Ca2+; ThermoFisher, cat: 14040141),

33.3μl 30mg/ml Collagenase (Sigma Aldrich, cat:C9697), 12.5μl 1% Trypsin (Sigma Aldrich, cat:T4174). For the collagenase digest method, ovaries were dissected out of young mice (21-25 days old) and rinsed in PBS. A small nick was made in each ovary using a fine needle. Each pair of ovaries were added to a vial of digestion mix and incubated at 37°C with constant agitation (550rpm) for 25 mins. After 5mins, the ovaries were pipetted up and down 5 times with a P1000 pipette, set to 50 0μl. 10 mins following, ovaries were pipetted gently, by up-taking the whole tissue with a P200 pipette, set to 200μl, around 40 times. 10 mins following, ovaries were pipetted 10 times with the P200 pipette, disrupting any clumps that remained in the ovary.

The digestion mix was then transferred to a 100mm coated dish (Nunc; Sigma Aldrich, cat:Z717223) and checked for viscosity under a dissection microscope. When most oocytes were free from somatic cells, and viscosity was not too high, 500 μl of M2 medium was added to the digestion mix to stop the digest. GV-stage oocytes were isolated from the mixture and washed through 4-5 droplets of M2 medium using a mouth pipette. Oocytes were rinsed once in PBS, and snap frozen on dry ice in 2-5 μl RLT+ buffer (QIAGEN, Germany, cat: 79216) in low-bind 0.2 ml PCR strip tubes(ThermoFisher, cat:AB-2000).

Embryo collections from mated females were performed during knockout characterisation, to determine the fertility of different genotype combinations in the *Nlrp5* mutant colony, and to investigate whether this knockout recapitulated the 2-cell arrest phenotype *in vivo*. Mated females were plug-checked to estimate timing of fertilisation, and oviducts and uteruses were dissected out at E3.5. Tissues were cleaned in M2 medium using tweezers and fine needles, trimming away fat. A syringe was filled with M2 medium, and inserted into one end of the oviduct, all within a large droplet of M2 medium. The liquid was expelled from the syringe gently, rinsing any embryos in the oviduct out into a glass collection dish. Embryos were gently rinsed in PBS using a mouth pipette, and visually staged under a dissection microscope.

#### Immunofluorescence staining of oocytes and embryos

Fixed oocytes or embryos were washed in PBS 0.05% Tween-20 (PBT), permeabilised in PBS 0.5% Triton X-100 for 1 hour at room temperature, and then washed again in PBT. Samples were blocked for 1 hour at room temperature, in PBS 0.05% Tween-20 1% BSA (BS) (BSA from Sigma Aldrich, cat: A9418), and then incubated for 1 hour at room temperature with the appropriate dilution of the primary antibody in BS (NALP5 rabbit polyclonal antibody gifted by Jurrien Dean, NIH, USA, 1:50. DNMT1 Abcam UK, cat:ab87654, 1:500. UHRF1, MBL Lifescience Japan, cat:D289-3, 1:100. DNMT3L, gifted by Shoji Tajima, Osaka University, Japan, 1:100.

α-tubulin, Invitrogen USA, cat:236-10501, 1:100. γ-tubulin, Invitrogen, cat:MA1- 19421, 1:100). Samples were then washed in BS for at least 1 hour at room temperature, and then incubated with the appropriate secondary antibody (AlexaFluor 488, 561 or 647, 1:500, ThermoFisher Scientific) and 2 drops of Hoechst reagent (NucBlue Live Ready Probes, 33342, Thermofisher, UK, cat:R37605) in 500μl BS for 1 hour at room temperature in the dark. Samples were washed for 1 hour at room temperature in PBT, protected from light.

#### Fluorescence Imaging and analysis

Imaging of NALP5-stainined oocytes was performed on an Andor Dragonfly600 spinning disk confocal microscope, at the Eugin Clinic’s Basic research laboratory, Parc Científic de Barcelona, Spain, in collaboration with Styliani Galatidou and Montserrat Barragan. All other immunofluorescence imaging was performed at the Babraham Institute imaging facility using a Zeiss LSM 780 confocal. Images were processed using ImageJ FIJI, to quantify fluorescence intensities. Fluorescence values were measured for the nucleus, cytoplasm, and background fluorescence for each oocyte image. Fluorescence plots were normalised by α or γ-tubulin fluorescence and/or background fluorescence. More details on data presentation and significance (p-values) can be found in the figure legends. For assessment of NSN-SN ratio differences, the proportion of oocytes in each stage was calculated for each mouse biological replicate, and boxplots were generated using ggplot2 (version 3.4.4). Significance was calculated using a t-test within the stat_compare_means function from the ggpubr (version 0.6.0) package (https://www.rdocumentation.org/packages/ggpubr/versions/0.6.0, version 0.6.0).

More details on data presentation and significance (p-values) can be found in the figure legends.

#### Time-lapse imaging of embryos

IVF for time-lapse imaging of developing embryos was performed at the BSU. Oocytes were collected from superovulated four to ten-week-old C57Bl/6 females. They were cleaned in M2 medium (Sigma Aldrich, MR-015P-5F) with added hyaluronidase (Sigma Aldrich, H2126) and then IVF was performed with C57Bl/6 sperm. Fertilised embryos were cultured in EmbryoMax KSOM medium (Merck, USA) under humidified conditions with 5% CO2 air composition at 37°C for 72 hours in the Geri+ timelapse incubator (Genea Biomedx, UK), and imaged at 5 minute intervals.

#### Mass spectrometry

Mass spectrometry analysis of GV oocyte samples was performed at the Babraham Institute Mass Spectrometry facility. 3-4 bulk oocyte samples per genotype (33 GV oocytes per bulk sample) were solubilised in SDS-PAGE loading buffer and run approximately 5mm into SDS-PAGE gels. After Coomassie-staining, gel pieces containing the entire protein-containing region from each lane were excised, and half of each band was destained, reduced, carbamidomethylated, and digested with trypsin. 20% of the resulting peptide digests were separated using an Ultimate 3000 nanoHPLC (ThermoFisher Scientific) on a reversed-phase column (0.075 x 500mm, Reprosil C18AQ, 3µm particles) with a 120min linear gradient from 2 to 35% acetonitrile, containing 0.1% formic acid, at a flow rate of 300nl/min. The column was interfaced to an Orbitrap Eclipse mass spectrometer (ThermoFisher Scientific) operating in data-independent acquisition mode. The mass spectrometric data were searched against the Uniprot canonical mouse proteome database (version July 2022) using DIA-NN software (v1.8.1)^74^, with further processing of the results in Perseus (https://maxquant.net/perseus/).

#### DNA and RNA isolation

DNA and RNA from individual oocytes were harvested from individual oocytes using the Smart-seq2-based G&T-seq protocol^75^.

#### Single-cell RNA sequencing library preparation

cDNA libraries (product of the G&T-seq protocol) were processed into scRNA- sequencing libraries using the Nextera XT DNA Library Preparation Kit (Illumina, FC- 131-1096) according to the manufacturer’s protocol for RNA-sequencing libraries.

Library concentrations were quantified on the Bioanalyzer (Agilent technologies, USA), and via KAPA Library Quantification Kit (KAPA Biosystems, USA). Both values were averaged, and the samples were pooled at a volume and concentration suitable for sequencing.

#### Single-cell PBAT sequencing library preparation

Single cell methylation analysis of oocytes was performed as previously described (Castillo-Fernandez et al. 2020), with 5 rounds of first strand synthesis. Library concentrations were quantified on the Bioanalyzer (Agilent technologies), and via KAPA Library Quantification Kit (KAPA Biosystems). Both values were averaged, and the samples were pooled at a volume and concentration suitable for sequencing.

#### scPBAT and scRNA sequencing

All scPBAT and scRNA-seq libraries were sequenced on an Illumina NextSeq500 High Output sequencer in 150bp paired end mode, at the Babraham Institute Genomics facility.

### QUANTIFICATION AND STATISTICAL ANALYSIS

#### Proteome analysis

Using Perseus software, the result from DI-AA for wild-type, *Nlrp5* heterozygous KO and *Nlrp5* homozygous KO samples were plotted using Principal Component Analysis (PCA) to determine if there was genotype-based clustering, then processed into a list of log-normalised protein abundances for each protein detected, per sample (Perseus standard pipeline, including t-test for comparison between genotype groups, with significance cutoffs off Benjamini-Hochberg adjusted p-value (FDR) < 0.1 and absolute log_2_ fold change > 0.5). Differentially abundant proteins were plotted on volcano plots. Significance cut-offs/parameters are detailed in the figure legends. Log-normalised protein abundance counts were plotted using the Heatmap function of the ComplexHeatmap R package (version 2.14.0; https://bioconductor.org/packages/release/bioc/html/ComplexHeatmap.html). First, all isoforms were plotted raw to visualise overall abundance of the isoform. Next, the per-isoform mean across samples was subtracted from each value to visualise the relative changes in abundance between samples. Where isoforms were subset, the heatmap calculation (and thus the clustering internal to the method) was performed on only the subset. Mass spectrometry datasets from 2 other KO studies were integrated into this analysis and compared with the *Nlrp5* KO proteomics data (*Nlrp14* KO^36^ and *Tle6* & *Padi6* KOs^3^). These datasets were chosen for comparative analysis due to either their putative or known membership of the SCMC. The relevant datasets were downloaded from the supplemental material of their associated papers, acquired from the PRIDE protein database, or acquired directly from the authors. Enrichment analyses were performed in R using gprofiler. Further comparative analysis between protein datasets were performed in R using the pcaMethods package (ppca method).

#### Sequencing data processing and quality control

Quality control and data processing were performed by the Bioinformatics team at the Babraham Institute. For single cell RNA-seq data processing, the raw data files were processed using the Babraham Institute Bioinformatics team’s standard nf_rnaseq pipeline (https://github.com/sandrews/nextflow_pipelines/blob/master/nf_rnaseq). This pipeline processes FastQ files, performs read count QC, contamination QC, quality-/adapter trimming using Trim Galore!, and splice-aware alignments to the GRCm39 mouse genome using HISAT2. Finally, it generates an aggregate QC report. For scPBAT sequencing data, the Babraham Institute nf_scBSseq pipeline was used for processing (https://github.com/s-andrews/nextflow_pipelines/blob/master/nf_bisulfite_scBSseq), with the additional parameters --trim_galore_args=“--clip_r1 9 --clip_r2 9” to account for potential mis-priming issues from the PBAT. The nf_scBSseq workflow runs an entire single-cell Bisulphite-seq (scBS-seq) processing pipeline on FastQ files. This includes QC, quality-/adapter trimming using Trim Galore!, contamination QC (post-trimming), alignments to the GRCm39 mouse genome using Bismark, deduplication, methylation extraction, and coverage file generation. Finally, it generates an aggregate MultiQC report. The workflow is usually carried out in ’--single_end’ mode, even for paired-end libraries (see https://felixkrueger.github.io/Bismark/faq/single_cell_pbat/).

#### Transcriptome and DNA methylation analysis

##### Analysis of scRNA-seq data

RNA-seq samples with fewer than 4 million reads were filtered out. The remaining samples were screened by the standard multi-QC report function of SeqMonk and determined to be of good quality (mitochondrial genes <2%, ribosomal rRNA <10%, 37-50% genes measured, >70% uniquely mapped reads in each sample). The oocyte samples were further subdivided into two maturity groups (Non-Surrounded Nucleolus (NSN) and Surrounded Nucleolus (SN)) based on the expression of NSN and SN-specific marker genes (Table S1(i)^5^). Samples that could not be clearly identified as either NSN or SN stage based on these marker genes were excluded, leaving 8 NSN and 8 SN samples to analyse (NSN=4 *Nlrp5* -/-, 4 *Nlrp5 +/+* (WT), SN= 2 *Nlrp5* -/-, 6 *Nlrp5 +/+* (WT)). Differentially expressed genes (DEGs) between WT and *Nlrp5* homozygous KOs were determined for each maturity group separately, using DESeq2 package in R^76^ (significance FDR <= 0.05; log_2_ FC >= 0.5). PCA plots were made for each maturity group separately, to indicate genotype-based clustering of samples. DESeq2 results were plotted as volcano plots, using the DESeq calculated adjusted p-value (Benjamini-Hochberg-corrected p-value) and DESeq2-calculated log_2_ fold change. All genes were plotted, and genes with an absolute log fold change greater than 2 and an FDR of less than 10% were plotted as significant (in red, and contributing to the numeric counts at the top of the volcano plot). The 10 most significant (lowest adjusted p-value) genes were additionally labelled (when there were at least 10 significant hits). DEGs from both NSN and SN groups were combined (total 584 unique DEGs) for the comparison with other omics datasets (proteomics/methylation). Gene enrichment analyses performed in R using gprofiler.

##### Analysis of scPBAT-data

Processed data were analysed in Seqmonk version 1.48.1. Processed methylation files were loaded into Seqmonk and methylation profiles for all libraries were compared to a wild-type oocyte methylome reference dataset (Castillo-Fernandez et al., 2020) using the genome viewer, to estimate coverage and rule out somatic cell DNA contamination (unlike somatic cells, oocytes have a characteristic methylation pattern of hyper and hypomethylated regions, with hypermethylated regions overlapping with transcriptional units). Quality control was performed on the scPBAT samples, filtering out samples with low read count (<600,000 CpGs covered) and samples with presumed somatic cell DNA contamination. Contaminated samples were inferred by 3 independent methods: 1) high methylation of X chromosome CpG islands (> 10%), which are hypomethylated in normal, non-contaminated oocytes; 2) modified proportion of the methylated cytosines in different contexts (taken from the Bismark QC output) compared to the normal ratio of CG <40%, CHG >3%, CHH>4% (Shirane et al., 2013); 3) modified histograms of the methylation of 100-CpG tiles compared to the standard bimodal profiles, peaking at 0 and 100% of methylation, with quantification over 100 CpG windows, using methylation pipeline with minimum of 5 calls per probe for individual oocytes. Only oocytes with satisfactory parameters to pass all 3 screening criteria were retained for further analysis. No *Nlrp5* +/- samples had sufficient coverage after MultiQC report assessment to proceed with downstream analyses. Between *Nlrp5 +/+* (WT) and *Nlrp5* -/- samples, there was no difference in the proportion of oocytes inferred as being contaminated.

The remaining WT and *Nlrp5* homozygous KO oocyte samples were pseudo-bulked in Seqmonk (10 WT and 19 *Nlrp5* -/- samples), resulting in a final CpG coverage of 28 and 31.6 million reads for WT and *Nlrp5* -/- groups, respectively. Analysis of the global DNA methylation was performed in SeqMonk, using initial quantification over 100 CpG tiles (minimum 30 calls per probe; Figure 7A), with further filtering of probes on different features: either contained within hypo/hyper-methylated domains (152,071 probes), or overlapping with hypermethylated CpG islands (> 75% methylation at least in one group, WT or *Nlrp5* homozygous KO; 1796 probes), or overlapping with gDMRs (germline differentially methylated regions) (61 probes)(Figure 7C). Identification of differentially methylated regions (2452 hits, overlapping with 1402 genes) amongst the probes contained within hypo/hyper- methylated domains was carried out using a Chi Square test with significance cutoffs of the adjusted P-value <0.05 and % methylation difference of >25% (Figure 7B).

The differential methylation of gDMRs and their heatmap were analysed using tiles directly over gDMRs (with quantification of minimum 20 calls per tile)(Figure 7D). The gDMRs were identified with Chi-square test with the above cutoffs on the significance. Only highly covered probes over differentially methylated gDMRs (>100 reads/1000bp) were considered as significant in Tables S3(ii)(iii).

#### Multi-omic analysis

To determine whether there was a significant overlap between scRNA-seq and proteomics hits (DEGs and differentially abundant proteins respectively) hyperheometric tests were performed (indicating p-value for overlaps) in R. Transcriptomic and proteomic log_2_ fold changes were plotted against one another.

## Acknowledgements

We would like to acknowledge the invaluable support of the Babraham Biological support unit (BSU) for animal maintenance and husbandry, and the excellent guidance and assistance from the Babraham Proteomics, Imaging, Sequencing and Bioinformatics facilities, with special thanks to Judith Webster, Simon Walker, and Megan Hamilton. We would like to also acknowledge Simon Andrews and Sarah Inglesfield from the Babraham bioinformatics facility for initial sequencing data processing and QC, and Jurrien Dean and Shoji Tajima, for providing MATER and DNMT3L antibodies respectively.

## Author Contributions

Conceptualization, Leah Nic Aodha and Gavin Kelsey. Knockout mouse design, Leah Nic Aodha and Asif Nakhuda with assistance from the Babraham Biological Support Unit (BSU). Investigation and data collection, Leah Nic Aodha with assistance from Styliani Galatidou, Edyta Walewska, Christian Belton, Antonio Galvão, Hanneke Okkenhaug at the Babraham Imaging facility, and Bill Mansfield at the BSU. Mass spectrometry and proteomics dataset mapping, David Oxley and Lu Yu at Babraham Mass-spectrometry facility. NSN-SN marker gene list provided by Soumen Khan. Formal analysis and data visualisation, Leah Nic Aodha, Alexandra Pokhilko, and Leah U Rosen. Supervision, Gavin Kelsey and Monserrat Barragán. Writing – Leah Nic Aodha, editing of initial drafts – Gavin Kelsey, final draft – Leah Nic Aodha, in consultation with all authors.

